# Stress-induced c-fos expression in the medial prefrontal cortex of male rats differentially involves the main glial cell phenotypes

**DOI:** 10.1101/2023.12.13.571548

**Authors:** Adriana Aguilar-Delgadillo, Fernando Cruz-Mendoza, Sonia Luquin-de Anda, Yaveth Ruvalcaba-Delgadillo, Fernando Jáuregui-Huerta

**Affiliations:** Neurosciences department, Health sciences center, University of Guadalajara, Guadalajara, México; Department of Physiology, School of Medicine, National Autonomous University of Mexico, Mexico City, Mexico

**Keywords:** c-fos, stress, prefrontal cortex, glia, immunofluorescense

## Abstract

Stress poses a challenge to the body’s equilibrium and triggers a series of responses that enable organisms to adapt to stressful stimuli. The prefrontal cortex (PFC), particularly in acute stress conditions, undergoes significant physiological changes to cope with the demands associated with cellular activation. The proto-oncogene c-fos and its protein product c-Fos have long been utilized to investigate the effects of external factors on the central nervous system (CNS). While c-Fos expression has traditionally been attributed to neurons, emerging evidence suggests its potential expression in glial cells. In this study, our main objective was to explore the expression of c-Fos in glial cells and examine how acute stress influences these activity patterns. We conducted our experiments on male Wistar rats, subjecting them to acute stress and sacrificing them two hours after the stressor initiation. Using double-labelling fluorescent immunohistochemistry targeting c-Fos, along with markers such as GFAP, Iba-1, Olig2, NG2, and NeuN, we analyzed a series of 35 μm brain slices obtained from the medial PFC. Our findings compellingly demonstrate that c-Fos expression extends beyond neurons and is present in astrocytes, oligodendrocytes, microglia, and NG2 cells—the diverse population of glial cells. Moreover, we observed distinct regulation of c-Fos expression in response to stress across the different phenotypes. These results emphasize the importance of considering glial cells and their perspective in studies investigating brain activity, highlighting c-Fos as a response marker in glial cells. By shedding light on the differential regulation of c-Fos expression in response to stress, our study also contributes to the understanding of glial cell involvement in stress-related processes.

## Introduction

Stress is a challenging condition that disrupts the homeostasis of the central nervous system (CNS) and activates behavioral and physiological responses to cope with the stressor **[1,2]**. Acute stress induces a rapid increase in neurotransmission, neuronal activation, and hormone release, ultimately culminating in profound alterations in the physiological parameters of the brain **[3]**. In the face of a stressor, distinct brain regions are activated, and among them, the prefrontal cortex (PFC) has emerged as a region of particular sensitivity to stress **[4]**. The PFC is involved in various cognitive functions, including attention **[5]**, decision-making **[6]**, and working memory **[7]**.

The assessment of cellular activation involved in the effects of stress on the prefrontal cortex (PFC) and other brain regions can be accomplished through the examination of c-Fos protein expression, which is encoded by the proto-oncogene c-fos. This immediate early gene (IEG) possesses the capacity to modulate the expression of target genes (TGs) **[8]**, acting as a component of various inducible enhancer pathways (CRE, SER, SIE) **[9]**. The low basal expression of c-Fos in the absence of stimulation renders it a reliable marker for quantifying neuronal activation in response to diverse stimuli **[10]**. Consequently, the detection of its nuclear protein has been extensively employed to investigate CNS activation and delineate neural pathways **[11,12]**. Experimental studies employing social stress **[13–15]**, restraint stress **[16]**, photoshock **[17]**, electric shock **[18]**, and exposure to environmental noise **[19,20]** have consistently demonstrated alterations in c-Fos protein activation. These findings underscore the potential of c-fos and its protein as valuable tools for investigating gene transcription in response to exogenous factors and for elucidating the neural networks implicated in stress responses.

Emerging research has brought attention to the involvement of glial cells in mediating brain activity, prompting the investigation of whether these cells are also subject to c-fos expression induced by stress **[21]**. Glial cells are no longer regarded solely as passive support cells but have been discovered to have crucial roles in modulating neural activity, information processing, and stress responses **[22]**. For instance, astrocytes have the ability to release neurotransmitters and regulate synaptic activity **[23]**, while microglia are involved in regulating inflammation and synaptic pruning **[24]**. Oligodendrocytes and oligodendrocyte precursor cells (OPCs), on the other hand, are responsible for myelination and play critical roles in enhancing neural transmission speed and efficiency **[25]**. These findings suggest that glial cells significantly contribute to brain function and activity.

While c-Fos expression has primarily been associated with neurons, emerging evidence suggests that glial cells, including astrocytes, oligodendrocytes, microglia, and NG2 cells, may also express this immediate early gene. Several studies utilizing various approaches, such as in vitro experiments, animal models of brain injury, administration of serum or stimulants, exposure to lipopolysaccharides, and drugs, have provided evidence supporting the expression of c-Fos in glial cells **[21]**. Despite these findings, the expression of c-Fos in glial cells under aversive conditions remains poorly understood. Therefore, the present study aims to investigate the expression of c-Fos in different cell types in the brain, with a particular focus on glial cells, in response to acute stress. The study seeks to examine these expression differences within the medial prefrontal cortex, given its high susceptibility to stress.

## 1 Materials and methods

### 2.1 Animals

Male Wistar rats, aged 90 days, were utilized for this study. The rats were housed in a controlled environment with a 12:12 light-dark cycle, maintained at a temperature of 22 ± 2°C, and a humidity level of 70%. Prior to exposure to the stressful stimulus, the rats had unrestricted access to balanced food and water. The animals were procured from the in-house breeding facility at the West Center for Biomedical Research in Guadalajara, Mexico. All experimental procedures were conducted in accordance with the guidelines outlined in the National Institutes of Health guide for the care and use of Laboratory animals (NIH Publications No. 8023, revised 1978) and approved by the Institutional Ethics Commission, CI. 068-2014.

### 2.2 Varied Acute Stress protocol

For the experimental group, four rats were subjected to a combination of two different stressors, with a 10-minute rest period between each stimulus. The first stressor involved restraint stress, where each rat was confined inside a PET plastic cylinder measuring 8x18 cm, equipped with 5mm holes at each end to ensure proper ventilation. This setup allowed the rat to breathe comfortably while restricting its movement. The second stressor employed was forced swimming, following a modified Porsolt paradigm [26]. The rats were individually placed in a plexiglass cylinder measuring 45 cm in height and 30 cm in diameter. They were then compelled to swim for a duration of 15 minutes, while maintaining the water temperature at a constant 10°C [27]. Meanwhile, the control group consisted of four rats that were not subjected to any stressors and were kept under standard laboratory conditions.

### 2.3 Intracardiac perfusion

Ninety minutes after exposure to the first stressful stimulus, the animals were administered a sublethal dose of sodium pentobarbital (60 mg/kg) via injection. Following the administration, the animals were perfused through the left ventricle with a 0.9% sodium chloride solution, followed by a fixative solution consisting of 3.8% paraformaldehyde in 0.1 M phosphate-buffered saline (PBS) at pH 7.4. The perfusion process ensured the proper fixation of the brain tissue. After perfusion, the brains were carefully extracted and left in post-fixation for a period of 2 hours. Subsequently, the post-fixed brains were sliced into coronal sections with a thickness of 35 μm using a Leica VT1000E vibratome. Slices were taken at intervals of every four cuts, resulting in sections that were 140 μm apart. An average of 10 sections per cellular marker were obtained for each brain rat in total. This process was carried out until the medial prefrontal cortex of the rat was completely covered. The stereotactic coordinates of Paxinos were employed as a reference guide during this procedure, specifically targeting the range of bregma +4.20 to +2.52 mm [28].

### 2.4 Fluorescence immunohistochemistry

Since c-Fos expression has been largely probed under immunohistochemistry methods, we used fluorescence immunohistochemistry to evaluate c-Fos co-expression among the different brain cells. Each brain was used to process the five cellular markers of interest for our study. The tissue sections were initially washed in tris buffered saline with 1% Tween (TBST) to remove any residual contaminants. Subsequently, blocking of non-specific binding sites was performed by incubating the sections in TBST containing 5% goat serum for a duration of 1 hour. The blocking step was carried out at 37°C for 30 minutes, followed by an additional 30 minutes at room temperature. To detect the expression of c-Fos, the sections were incubated with anti-Fos antibodies (cell signalling (E8) and Santa Cruz (SGIABO253)), diluted 1:500 in TBST. The sections were incubated with the primary antibodies for a total of 54 hours at 4°C. For double labeling, additional antibodies were employed to detect specific cell types. These included mouse anti-NeuN (Millipore MAB377) at a dilution of 1:500, mouse anti-NG2 (Millipore 05-710) at a dilution of 1:100, rabbit anti-GFAP (Dako Z0334) at a dilution of 1:750, rabbit anti-Iba1 (Thermo Fisher 016-24461) at a dilution of 1:250, and rabbit anti-Olig2 (Millipore AB9610) at a dilution of 1:300. After 24 hours of incubation with the primary antibodies, the sections were rinsed in TBST and subsequently incubated in secondary antibodies. The secondary antibodies used were anti-rabbit Alexa and anti-mouse Alexa, both at a dilution of 1:1000 in TBST. The incubation with the secondary antibodies was carried out for 2 hours at a temperature of 20-25°C. The secondary antibodies used were provided by Invitrogen and were labelled with Alexa 488 and Alexa 594, respectively. Following the incubation with the secondary antibodies, the sections were rinsed in tris buffered saline (TBS) to remove any unbound antibodies and other residues.

### 2.5 Image processing

The sections were carefully mounted onto slides and subsequently examined under a Leica DMi8 inverted microscope. In order to ensure representative sampling, a systematic random approach was employed, with a sampling interval of 5. The sections of the medial prefrontal cortex (mPFC) were further subdivided into specific regions, namely the dorsal Anterior Cingulate Cortex (ACC), Prelimbic (PL), and Infralimbic (IL) cortex. Microscopic analysis was performed at a magnification of 20X, which allowed for the visualization of an area measuring 622.18 x 466.55 μm^2^. Two microscopic fields were considered per subregion of mPFC of each hemisphere. To aid in the analysis, the captured microphotographs were processed using the freely available software ImageJ. This software facilitated the manipulation and enhancement of the images, enabling a clearer visualization of the markers of interest. A manual quantification of the c-Fos+ cells and double labeling with neuron and glial cells was performed based on the processed images. Double labelled cells were corroborated on higher magnification (40x or 63x). To stablish the co-localization criteria, a Leica TCS SP2 confocal system was used. 50% cell overlap (c-fos + glial or neuron marker, among 3 z-stacks (0.9 μm each) was established as a minimum colocalization criteria. The analysis was performed for 2 different investigators blinded to treatment. Figure 1 provides a visual representation of the regions of interest, the specific markers being analysed, and the sampling method employed.

**Figure 1.**
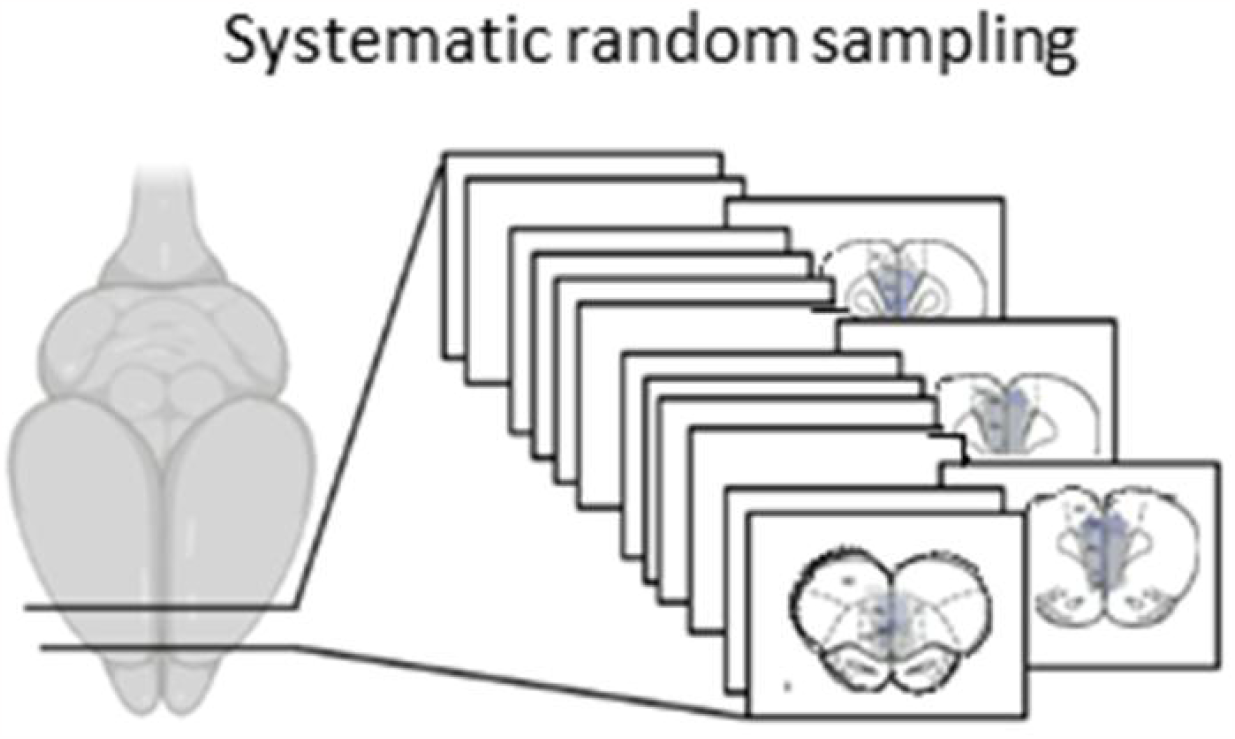
Provides a visual representation of the the systematic random sampling method used for cell counting.

### 2.6 Statistical analysis

Descriptive statistical analysis was employed to examine the data, including measures of central tendency (mean) and dispersion (standard error of the mean SEM). These measures provided a summary of the quantitative variable under investigation. To compare the differences between the stress and non-stress conditions in every cell types, student’
ss t test were conducted. One-way analysis of variance (ANOVA) was performed to further explore and compare differences between the lineages. To analyse expression differences between brain cell types after exposure to the stimuli we conducted a chi square test. As a complement, we conducted a mixed two-way ANOVA analysis to estimate the effect of factors implied on the experiment (stimuli and cell type) and its interaction. The impact of the effect was measured as partial eta squared and statistical significance was determined based on a predetermined threshold, typically set at p < 0.05.

## 2 Results

### 3.1 Acute stress increases c-Fos expression in mPFC

The number of c-Fos-positive cells was quantified in the Anterior Cingulate Cortex (ACC), Prelimbic Cortex (PL), and Infralimbic Cortex (IL) (Figure 2A) of both the control and acute stress groups. The analysis revealed a significant increase in c-Fos expression in the mPFC of the stress group compared to the control group (Figure 2B; p < 0.001). This enhanced expression was observed across all subregions of the mPFC, including the ACC, PL, and IL. Interestingly, there were no statistically significant differences detected between the subregions of the mPFC, indicating a similar level of c-Fos expression among these areas.

**Figure 2.**
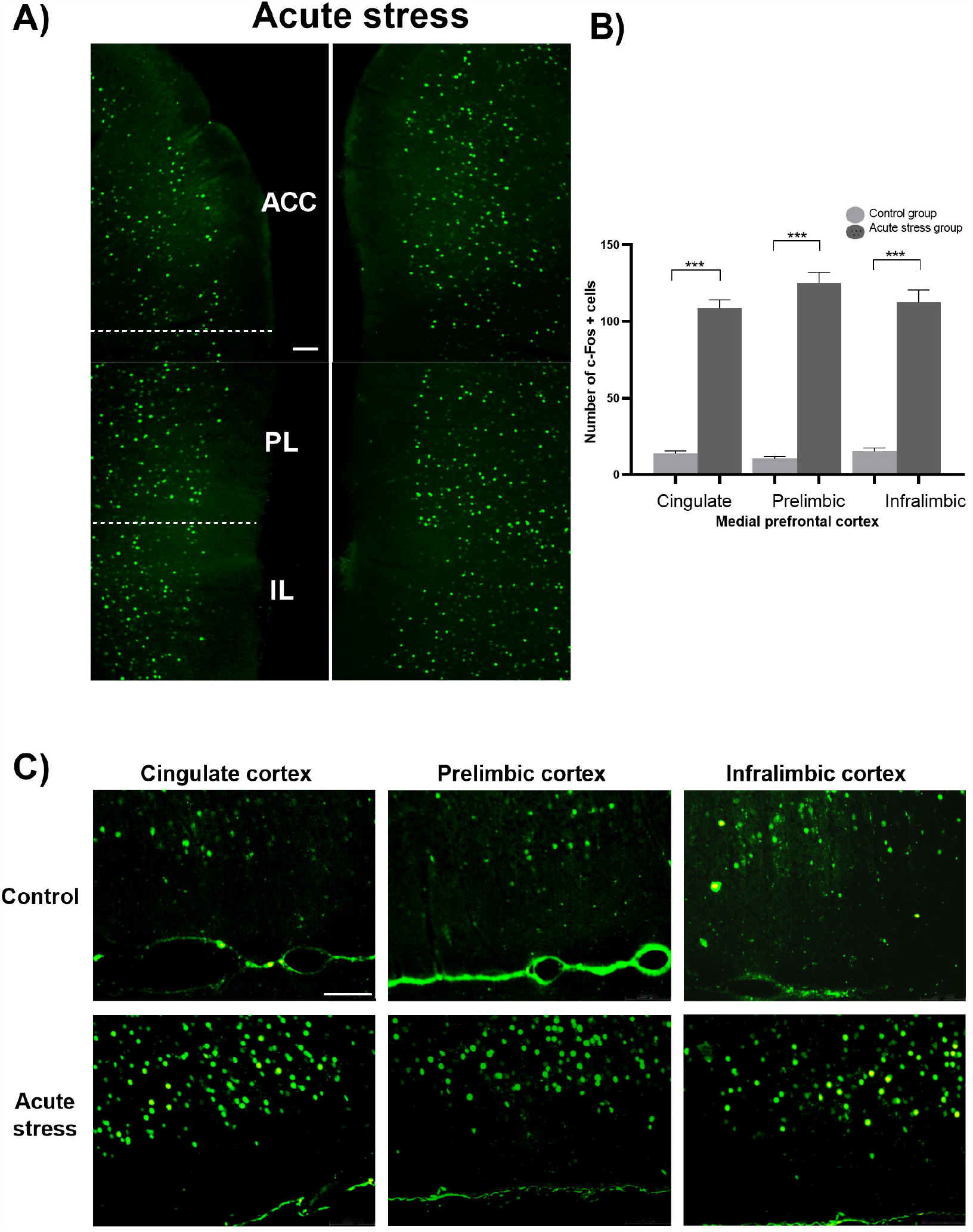
c-Fos expression in medial prefrontal cortex of rats exposed to acute stress. A) the figure displays the representation of c-Fos expression in the ACC, PL and IL at a magnification of 10X C) micrographs at a magnification 20X depicts the expression of c-Fos in control and stress groups in every subregion of the mPFC. Fluorescent micrograph has a scale bar 100 μm. B) the graph corresponding to the count of Fos-positive cells in the control and acute stress groups is presented. The data represents the mean ±SEM (One-way ANOVA; ***p < 0,001).

### 3.2 All evaluated cell types expressed the c-Fos proto-oncogene in mPFC

Furthermore, we conducted double labelling experiments to prove the co-localization of c-Fos with both neurons and glial cells. This action evidenced the presence of c-Fos in Astrocytes, Microglia, Oligodendrocytes, NG2 cells and neurons. Figure 3 illustrates the five evaluated lineages and demonstrates the presence of c-Fos in each phenotype.

**Figure 3.**
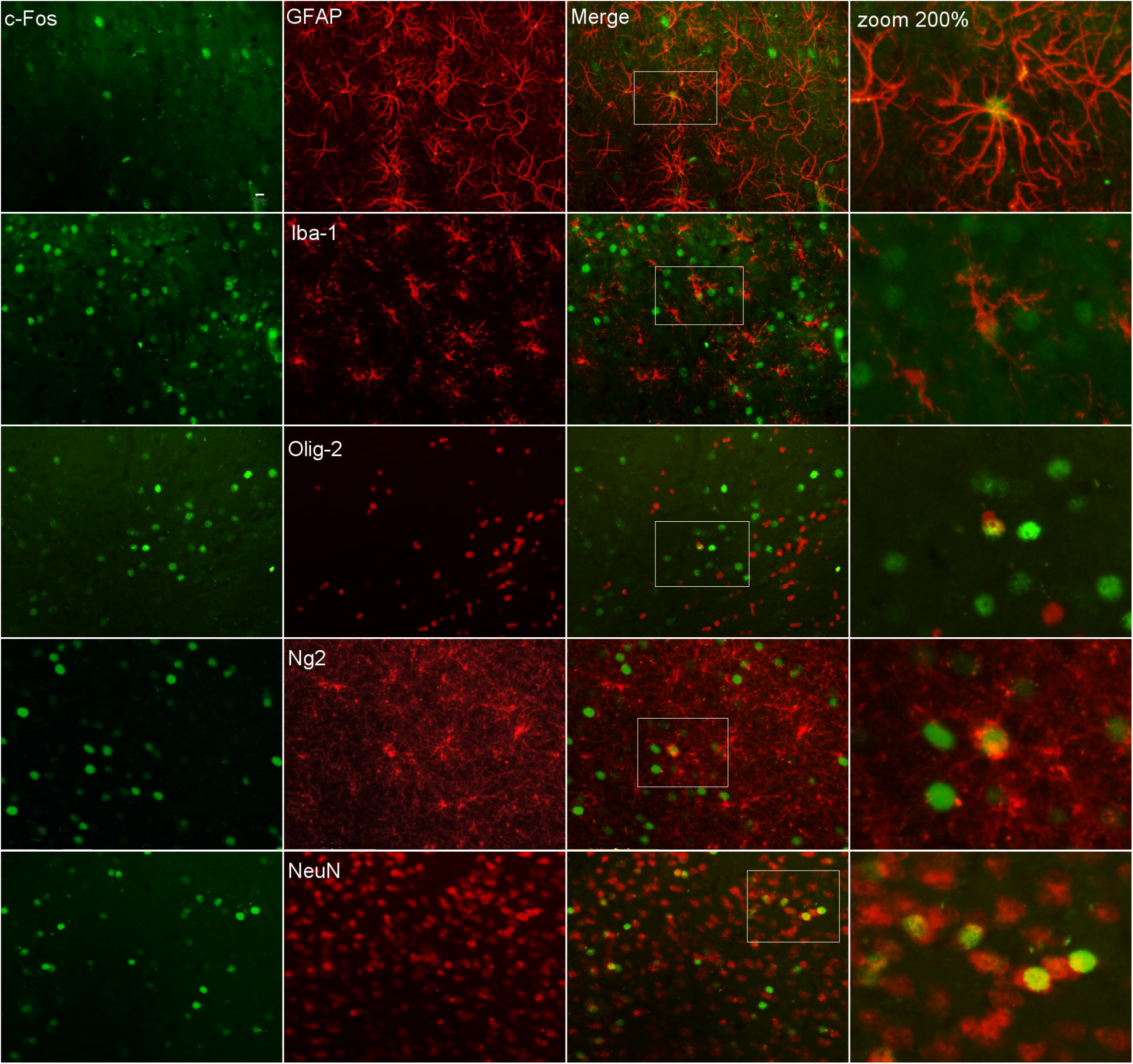
c-Fos expression in glial cells. The images show evidence of double labeling in both neurons and glial cells with c-Fos in the brains of rats exposed to acute stress. Green signal correspond to positive c-Fos cells (first column) and the red signal (second column) indicates positive cell labeling for glia (GFAP, Iba-1, Olig2, NG2) and neurons. The third column show merges at 40X magnification. Scale bar 10 μm. The white rectangles indicate the area that was zoomed in by 200% to illustrate a double labeling, which can be seen in the fourth column.

### 3.3 Acute stress differentially affected neurons and glial cells in mPFC

Next, we analysed the stress activated cells to stablish any possible difference associated to region and/or phenotype. Statistical analysis revealed significant differences between the stressed and control groups in the Anterior Cingulate Cortex (ACC) (Figure 4B; p < 0.001), Prelimbic Cortex (PL) (Figure 5B; p < 0.001), and Infralimbic Cortex (IL) (Figure 6B; p < 0.001).

**Figure 4.**
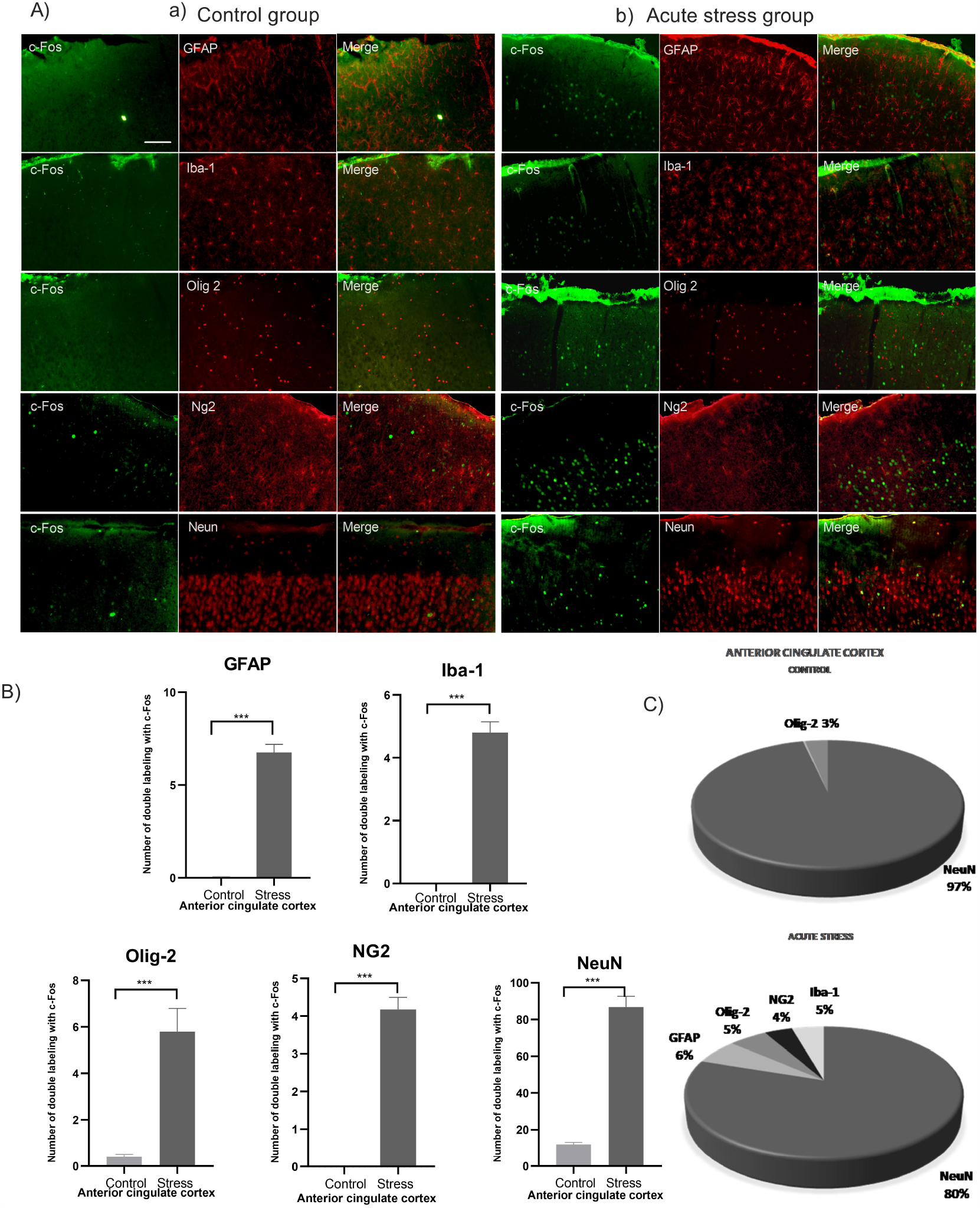
c-Fos expression by cell type in the Anterior Cingulate Cortex of rats exposed to acute stress. A) The images show the fluorescent micrographs of a) control and b) acute stress conditions. The green signal indicates the c-Fos+ cells, the red signal indicates lineage cell+ (GFAP, Iba-1, Olig2, NG2, and NeuN), and the third column shows the corresponding merge. Magnification 20X, scale bar 100 μm. B) Illustrates the count of double labeling of c-Fos/brain cell types in the control and acute stress groups. The data represents the mean ±SEM. (Student’
ss t test ; ***p < 0,001.C). The proportions of double labeling of c-Fos/neurons and c-Fos/glia are illustrated.

**Figure 5.**
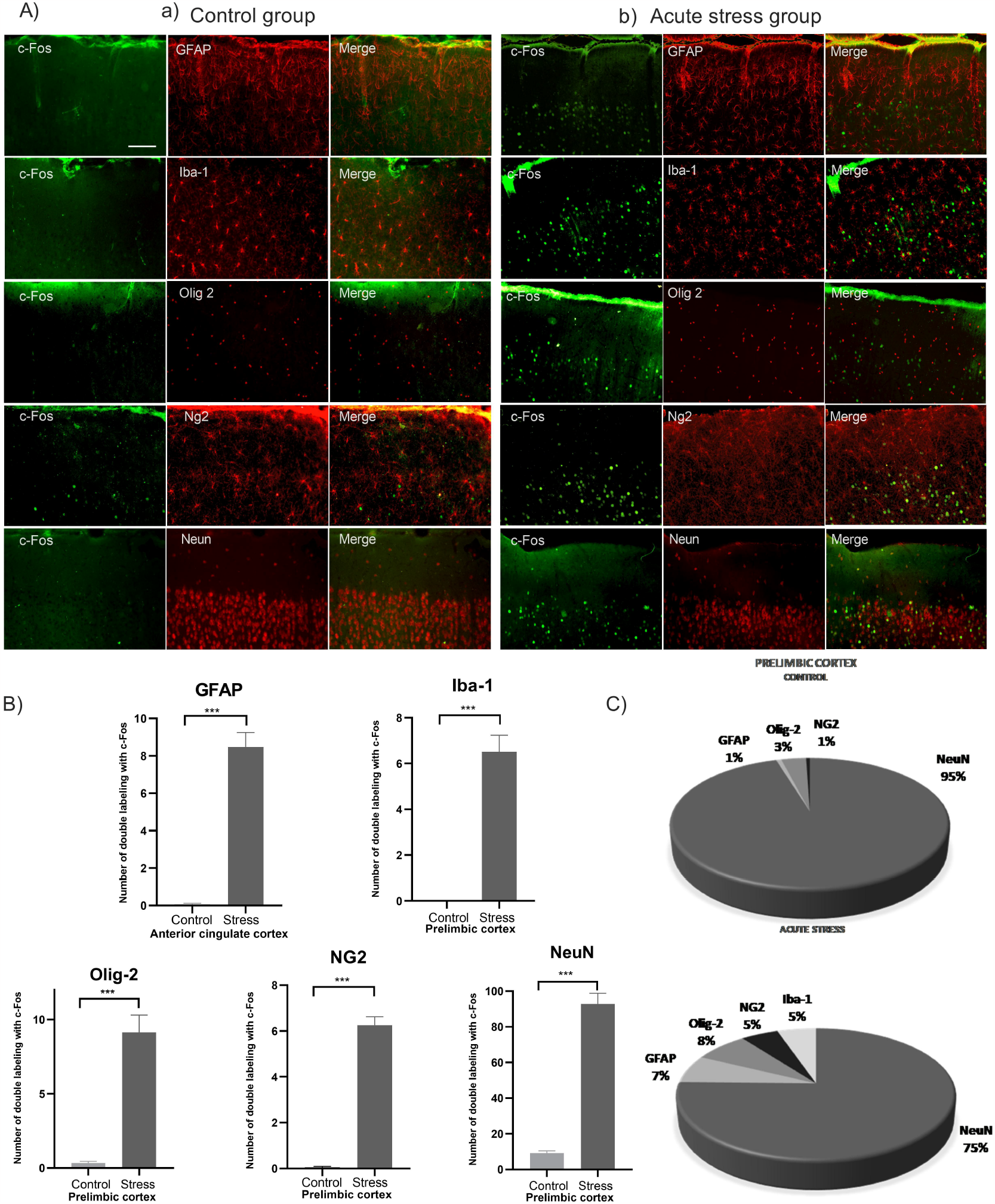
c-Fos expression by cell in the Prelimbic cortex of rats exposed to acute stress. A) The images show the fluorescent micrographs of a) control and b) acute stress conditions. The green signal indicates the c-Fos+ cells, the red signal indicates lineage cell+ (GFAP, Iba-1, Olig2, NG2, and NeuN), and the third column shows the corresponding merge. Magnification 20X, scale bar 100 μm. B) The graph shows the count of double labeling of c-Fos/brain cell types in the control and acute stress groups. The data represents the mean ±SEM. (Student’
ss t test ; *p < 0,05, **p < 0,01, ***p < 0,001). C) The proportions of double labeling of c-Fos/neurons and c-Fos/glia are illustrated.

**Figure 6.**
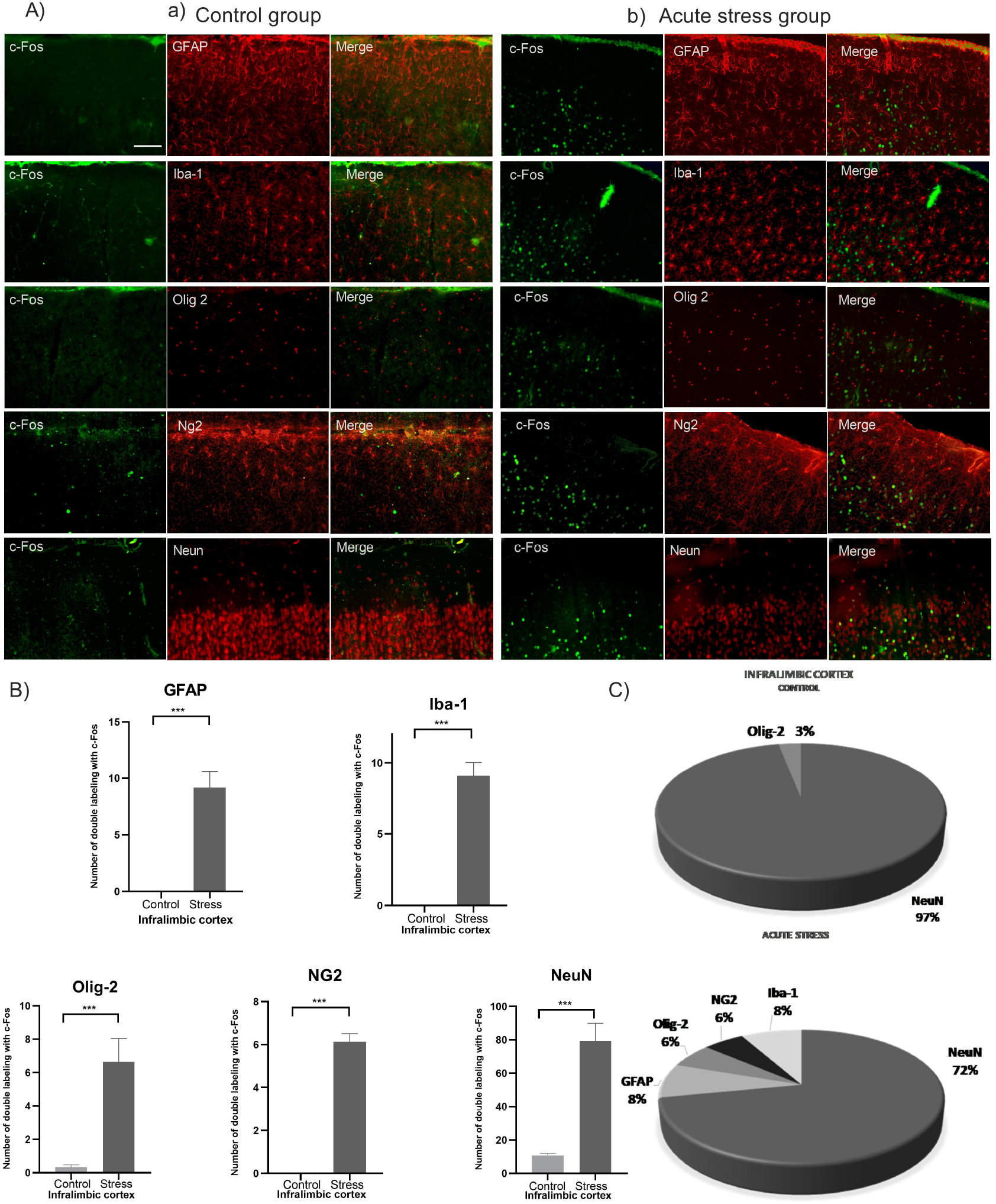
c-Fos expression by cell type in the Infralimbic cortex of rats exposed to acute stress. A) The images show the fluorescent micrographs of a) control and b) acute stress conditions. The green signal indicates the c-Fos+ cells, the red signal indicates lineage cell+ (GFAP, Iba-1, Olig2, NG2, and NeuN), and the third column shows the corresponding merge. Magnification 20X, scale bar 100 μm. B) The graph shows the mean of double labeling of c-Fos/brain cell types. The data represents the mean ±SEM. *p < 0,05 (Student’
ss t test; **p < 0,01, ***p < 0,001). C)The proportions of double labeling of c-Fos/neurons and c-Fos/glia are illustrated.

In the Anterior Cingulate Cortex, the analysis showed that 97% of the double-labeled cells in control group, corresponded to c-Fos/neuron, while only 3% were identified as c-Fos/glia. However, in the stress group, 80% of the labeled cells were c-Fos/neuron, and 20% were identified as c-Fos/glia. This shift in proportion was confirmed by a chi square test (χ^2^=11.89,4 p=0.0182).

In the Prelimbic Cortex, our analysis revealed that 95% of the double-labeled cells in control group were identified as c-Fos/neuron, and only 5% were classified as c-Fos/glia. However, under stress conditions, the proportions shifted to75% of the labeled cells being c-Fos/neuron, and 25% being identified as c-Fos/glia (χ^2^=13.57,4 p=0.0088).

In the Infralimbic Cortex, the control rats exhibited a predominance of c-Fos/NeuN cells, with 97% of the double-labeled cells being identified as c-Fos/NeuN and only 3% as glia. However, in the acute stress group, the proportions shifted, with 72% of the labeled cells being c-Fos/NeuN and a notable increase to 28% as c-Fos/glia (χ^2^=20.54,4 p=0.0004).

Then, we compared subregions of the mPFC and observed a higher expression of c-Fos/microglia-positive cells in the Infralimbic Cortex (IL) compared to both the Prelimbic Cortex (PL) and Anterior Cingulate Cortex (ACC). This indicates that microglia in the IL region may be particularly responsive to the acute stress stimulus compared to the other subregions of the medial prefrontal cortex (Figure 7A). Furthermore, we also found a higher expression of c-Fos/NG2-positive cells in both the Prelimbic and Infralimbic cortices compared to the ACC. This shows that NG2 cells may exhibit a greater response in these regions (Figure 7B), compared to the other glial types (figure 7C-E).

**Figure 7.**
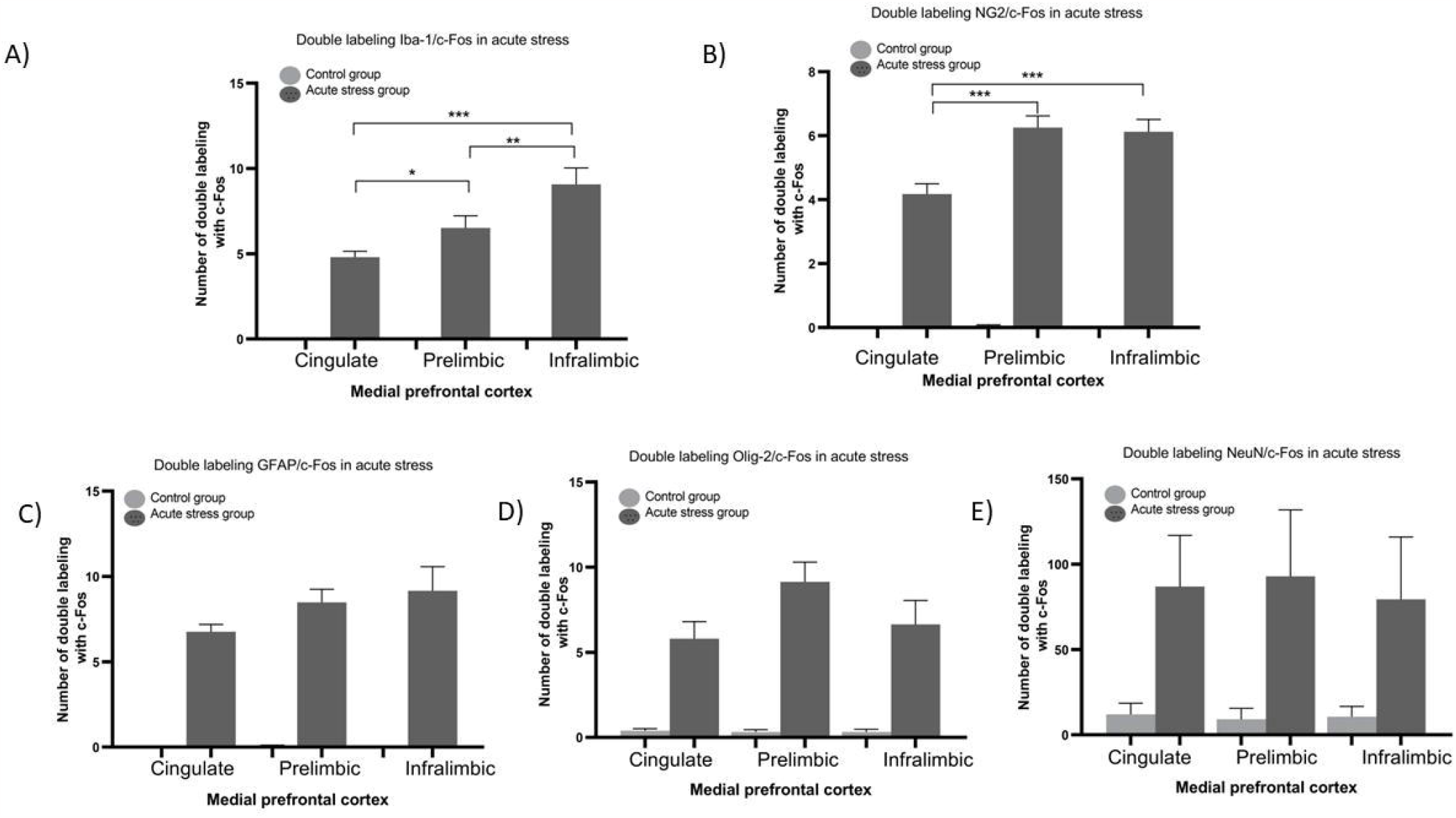
c-Fos expression in rats exposed to stress across subregions of mPFC. The graph shows the differences of double-labeled c-Fos/cell types between subregions of the medial prefrontal cortex of the rats exposed to acute stress. The data represents the mean ±SEM (One-way ANOVA; *p < 0,05 **p < 0,01, ***p < 0,001).

Finally, we measured the impact of the effect for the factors involved in our experiment. We found statistical significance for all subregions examined: ACC (Stimuli p= <0.0001, partial eta squared =14.23; Cell type p= <0.0001, partial eta squared=51.86 and an interaction effect p= <0.0001, partial eta squared =28.82), PL (Stimuli p= <0.0001, partial eta squared =9.992; Cell type p= <0.0001, partial eta squared=30.24 and an interaction effect p= <0.0001, partial eta squared =19.75) and for IL (Stimuli p= <0.0001, partial eta squared =18.14; Cell type p= <0.0001, partial eta squared=52.76 and an interaction effect p= <0.0001, partial eta squared =29.12). These results confirmed significant effect for both factors with a strong interaction between them.

## 4 Discussion

Our study provided evidence of the heightened sensitivity of the medial prefrontal cortex (mPFC) to acute stress, as indicated by a significant increase in c-Fos expression in animals exposed to stress. Notably, our findings went beyond neurons and revealed that glial cells also exhibited the expression of c-Fos, indicating their involvement in the stress response within this region. Furthermore, our data demonstrated that acute stress enhanced the proportion of glial cells capable of expressing this activity marker.

### 4.1 -c-Fos expression on the medial prefrontal cortex of stressed rats

It is known that transcriptional activation of the c-Fos gene in the brain is closely associated with neuronal depolarization. This process is influenced by the activation of specific channels, namely for example NMDA and VSCCs (voltage-sensitive calcium channels), which initiate signaling pathways that target response elements on the c-fos promoter. These response elements include the serum response element (SRE), cAMP response element (CRE), and SIF-inducible element (SIE) [29]. Under stressful conditions, additional signalling pathways associated with the NMDA receptor, such as the CREB/CRE and erk/MAPK/SER pathways, are also implicated in the upregulation of c-fos expression [30–33]. These pathways further contribute to the neuronal excitation and depolarization observed in response to stress. The upregulation of c-fos expression following extra-synaptic stimulation has been found to be more reliable and robust compared to other immediate early genes, such as Zif268 [34]. Therefore, c-Fos has become a widely used marker for studying neuronal activation and mapping neural circuits in response to various stimuli, including stress [18,35]. By utilizing c-Fos as an activation marker, researchers can gain valuable insights into the patterns of neuronal activation and the underlying neural circuits involved in stress responses.

This knowledge helps to deepen our understanding of the complex mechanisms through which stress affects the brain and provides a basis for investigating the functional consequences of stress-related neuronal activation. Then, findings of this study align with previous research indicating that the Anterior Cingulate Cortex (ACC), Prelimbic Cortex (PL), and Infralimbic Cortex (IL) are activated in response to acute stress protocols. The specific activation patterns observed in different studies may vary depending on the intensity and nature of the stressors employed [36]. In this study, using a combined acute stress protocol, no statistically significant differences were found in c-Fos expression between the subregions of the medial prefrontal cortex (mPFC). Similar results have been reported for the acquisition of contextual fear responses [37], stimulus-response associations [38], assessment of controllability, behavioural and vegetative alert responses [39], cardiovascular adjustments [40], and control of avoidance and extinction responses [41]. The absence of significant differences between the subregions of the mPFC implies that they may function collectively to orchestrate various aspects of the stress response. This coordinated activation within the mPFC subregions likely contributes to the integration of emotional, cognitive, and physiological responses that occur during stressful situations.

### 4.2 Expression of c-Fos in glia: possible roles and mechanisms associated with stress

Our findings reveal that stress can induce the expression of c-Fos in non-neuronal cells of the brain, which do not typically undergo depolarization. This raises intriguing questions about the activation mechanisms and the potential role of c-Fos in glial cells under stress. While the effects of acute and chronic stress on neurons and glia have been extensively studied, the scientific literature on the specific study of glial cells in the context of acute stress is limited. Nonetheless, it is well-established that acute stress does impact glial cells, although their involvement in the stress response is not as well understood as that of neurons. Prior to our study, the role of c-Fos in glia under stressful conditions had not been thoroughly investigated. Existing studies primarily focus on the relationship between Fos and glia in injury and disease models, often involving non-murine species [21].

#### 4.2.1 Roles of c-fos in glia under stress conditions

Studies have shown that astrocytes, undergo various changes in response to stress. These changes involve alterations in transcriptional regulation associated to modifications in the morphology and establishment of contacts between astrocytes [42,43]. Additionally, stress has been found to increase the release of extracellular gliotransmitters and growth factors by astrocytes, influencing intercellular communication [44,45]. Furthermore, stress can also impact astrocyte density, potentially leading to changes in their overall population within the brain [46,47]. Microglia, also undergo modulation in response to acute stress. These changes involve alterations in their activation state [48,49], morphological transformations [49–51], sensitization to subsequent proinflammatory stimuli [52,53], modulation of anti-inflammatory responses [49], dose-dependent increases in proliferation [54,55], and modifications in transcriptional regulation [52]. Then, it is clear that acute stress can influence proliferation, morphology, and functional characteristics of astrocytes and microglia. Studies have indicated that the c-fos gene may play a role in regulating proinflammatory responses and mediating morphological and functional alterations in glia [56,57]. Therefore, it can be hypothesized that c-fos is involved in the modulation of morphological, proliferative, and functional changes associated with the glial response to acute stress.

Most studies examining the effects of stress on oligodendrocytes and NG2 cells have primarily focused on chronic stimulation. There is evidence linking glucocorticoids to the initiation and enhancement of myelin formation [58,59], increases in oligodendrogenesis [60], and higher cell density [61]. Also, NG2 cells have shown changes in density [62] and cellular activation [63] in response to chronic stress. While fewer studies have specifically investigated the acute responses of oligodendrocytes and NG2 cells to stress, previous research has examined the effects of acute neuron activation on oligodendrocyte precursor cells (OPCs). Optogenetic stimulation of neurons has been shown to induce a proliferative response in OPCs and increase myelin thickness [64]. This suggests that neuronal activation can influence the proliferation, differentiation, and myelination of OPCs, thereby affecting cellular plasticity. Recent studies confirmed that c-fos, along with c-jun, may contribute to the proliferation and differentiation towards oligodendrocytes [65]. These studies proposed PKC as a potential mediator of these processes [65,66]. Furthermore, c-Fos has been involved in the differentiation of oligodendrocyte progenitor cells (OPCs) into mature oligodendrocytes [67]. Notably, the induction of c-Fos has been observed in oligodendrocyte progenitors stimulated by norepinephrine [68]. It has been also reported that c-Fos expresses in oligodendrocytes after excitatory stimulation [69]. Hence, it seems plausible that c-Fos also regulate proliferation, differentiation, and maturation of these cell lineages under stressful conditions.

#### 4.2.2 Mechanisms associated with the induction of c-fos in glia

The mechanisms underlying the induction of c-fos in glial cells remain unknown. While in neurons, c-fos is primarily induced by depolarization and changes in calcium concentration, serving as a marker of synchronous activation, similar mechanisms cannot be attributed to glia as they do not generate action potentials. Therefore, it would be valuable to explore in greater detail the specific mechanisms by which c-fos is induced in glial cells under conditions such as acute and chronic stress. One potential avenue to investigate is the integrative mechanisms involved in glia-to-glia and glia-to-neuron communication. For instance, the release of glutamate in excitatory synapses could activate glutamate receptors in glial cells, subsequently initiating signalling pathways that lead to the transcriptional activation of genes like c-fos. This hypothesis finds support in studies that have reported the induction of c-fos by glutamate in astrocytes through NMDA receptors [70], oligodendrocyte progenitors through AMPA-R and KA-R receptors [71], and microglia through ionotropic and metabotropic receptors [72]. In the case of microglia, their activation could be associated with the mitogen-activated protein kinases (MAPKs) pathways [57]. Another possibility to consider is the direct regulation of the c-fos gene by glucocorticoids through their interaction with glucocorticoid receptors (GR) in glial cells. This could involve the activation of various signalling pathways, as previous studies have reported the joint involvement of GR and NMDA receptor-Erk-MAPK pathways in the transcriptional regulation of genes like c-fos in neurons following psychological stress [73,74]. The direct regulation of c-fos by glucocorticoids in glial cells could imply the engagement of similar pathways. Regarding the results presented in our study on the prefrontal cortex, the increased expression of c-Fos observed after acute stress could be associated, in the initial stages, with the negative feedback response of the medial prefrontal cortex (mPFC) to the initial elevation of glucocorticoids induced by acute stimulation [75]. Neuronal activation in the mPFC may be attributed to an increase in excitatory synaptic strength [76], while other neurotransmitters such as dopamine [77] and norepinephrine [78] could also be involved. These neurotransmitter systems likely contribute to the complex interplay of factors that influence c-Fos expression in the mPFC under acute stress conditions.

It is important to note that these proposed mechanisms are based on our understanding of neuronal processes. However, it is plausible that the transcriptional induction of c-fos in glial cells occurs through distinct mechanisms and time dynamics. Therefore, further research is necessary to explore and elucidate the specific mechanisms by which c-fos transcription is induced in glial cells.

### 4.3 Increased proportion of glial cells capable of expressing c-Fos in mPFC under acute stress

Our study evidenced the concurrent activation of glial cells in the medial prefrontal cortex (mPFC), which exhibited minimal c-Fos expression under control conditions but presented a significant 20 to 30% increase in response to acute stress. This increase in glial c-Fos expression paralleled the observed neuronal activation, suggesting that glial cells may play a crucial role in meeting the demands of activated neurons. Specifically, we found statistically significant differences in the co-expression of c-fos in microglial cells within the prelimbic (PL) and infralimbic (IL) cortices. This suggests that microglia may directly contribute to the neuronal responses and alterations associated with mPFC disfunction, as has been previously reported under other stress models [79]. Furthermore, it is worth noting that microglia may become sensitized to future stimuli, potentially influencing the susceptibility of the mPFC cortex to pathological conditions [80].

Similarly, we also observed a higher expression of c-Fos in NG2 cells in the PL and IL regions. This increased expression could be associated with transient elevations in activation, and the fact that NG2 cells are among the first glial cells to respond to damage and stress [62]. Both the IL and PL cortices may exhibit more sensitive changes in their immediate response to stressors, thus requesting a greater involvement of specific glial cell populations. Overall, our findings shed light on the coordinated activation of glial cells within the mPFC under stress conditions, highlighting their potential contribution to neuronal responses and adaptive processes. These findings provide valuable insights into the complex interplay between neurons and glial cells in the mPFC and emphasize the importance of studying glial responses in the context of stress and related disorders.

## 3 Conclusions and perspectives

c-Fos has traditionally been used as a marker of neuronal activation in response to various stimuli, including stress. Its involvement in processes such as proliferation, transformation, differentiation, and cell death has been well-documented in neurons. However, the role of c-Fos in glial cells has remained elusive. This groundbreaking study represents the first investigation exploring c-Fos expression in glial cells. By exploring the mechanisms associated with non-neuronal cell activation, this study provides new insights and perspectives on the potential use of c-Fos as a marker of activation and plasticity/transformation. While this represents an important initial step in understanding the involvement of c-Fos in glia under stress, there are still numerous aspects to be explored and confirmed. Further investigations are warranted to elucidate the mechanisms underlying gene induction, the regional and temporal dynamics of activation, and the specific roles of c-Fos in the acute and chronic stress responses of glial cells. Overall, this study pioneers the exploration of c-Fos as a glial plasticity/transformation marker, opening up new avenues for research and paving the way for a deeper understanding of the intricate interplay between glial cells and stress-related processes.

## Conflict of Interest

The authors declare that the research was conducted in the absence of any commercial or financial relationships that could be construed as a potential conflict of interest.

## Author Contributions

Conceptualization, F.J.-H. and S.L.; methodology, A.A.-D. and F.C.-M.; software, A.A.-D. and F.C.-M.; validation, F.J.-H., Y.R.D. and S.L.; formal analysis, A.A.-D., F.J.-H. and F.C.-M.; investigation, F.C.-M. and F.J.-H.; resources, S.L., Y.R.D. and F.J.-H.; writing—original draft preparation, A.A-D., F.C.-M., F.J.-H. and Y.R.D.; writing—review and editing, F.J.-H., Y.R.D., A.A.-D., F.C.-M and S.L.; supervision, F.J.H, and S.L.; project administration, F.J.-H. and Y.R.D.; funding acquisition, F.J.-H. and Y.R.D. All authors have read and agreed to the published version of the manuscript.

## Funding

This work was supported by the “Consejo Nacional de Ciencia y Tecnología” (CONACyT México), grant numbers CF-2023-G-243 for Limei Zhang Ji, 238313 for YRD, 221092 for FJH, and Programa de Fortalecimiento de Institutos, Centros y Laboratorios de Investigación, CUCS, UdG 2023.

## Acknowledgments

We thank the Biomedical Sciences program at the University of Guadalajara for supporting our students’ activity in this project. We also thank the Department of Neurosciences for the provisions made to our lab.

## Notes

### Competing Interest Statement

The authors have declared no competing interest.

